# Boosting the localization precision of dSTORM by biocompatible metal-dielectric coated glass coverslips

**DOI:** 10.1101/136739

**Authors:** Hannah S. Heil, Benjamin Schreiber, Monika Emmerling, Sven Hoefling, Martin Kamp, Markus Sauer, Katrin G. Heinze

**Affiliations:** Rudolf Virchow Center, Research Center for Experimental Biomedicine, University of Würzburg, Josef-Schneider-Str.2, 97080 Würzburg, Germany; Technische Physik, Physikalisches Institut and Wilhelm Conrad Röntgen-Center for Complex Material Systems, University of Würzburg, Am Hubland, 97074 Würzburg, Germany; SUPA, School of Physics and Astronomy, University of St Andrews, St Andrews, KY16 9SS, United Kingdom; Department of Biotechnology and Biophysics, Biozentrum, University of Würzburg, Am Hubland, 97074 Würzburg, Germany

## Abstract

Super-resolution techniques such as direct Stochastic Optical Reconstruction Microscopy (dSTORM) have become versatile and well-established tools for biological imaging over the last century. Here, we theoretically and experimentally show that clever combination of different fluorescence modalities allows further improvements. We found that the interaction of fluorophores with plasmonic surfaces boost super-resolution performance in dSTORM approaches as it allows for tailoring the excitation and emission properties. The strength of the approach is that no further specialized microscope setup is required as the described enhancement solely rely on metal-dielectric coated glass coverslips that are straightforward to fabricate. Such biocompatible plasmonic nanolayers enhance the signal-to-noise ratio of dSTORM, and thus sharpens the localization precision by a factor of two.

---

To date, advanced fluorescence microscopy methods for resolution enhancement are based on either on-off fluorophores (for PALM [1], STORM [2], dSTORM [3],[4] or STED [5]), Structured Illumination Microscopy (SIM) [6], 4Pi-microscopy [7] and deconvolution routines [8]. Despite their great success, all of these techniques have their advantages and challenges. Intrinsically, localization techniques usually only provide limited temporal resolution, but excellent localization precision in an affordable, straightforward setup. SIM together with deconvolution routines only provides a limited spatial resolution improvement and the need of expensive hard- and software, however lately provides quite impressive acquisition speeds ([9],[10]) for imaging of tick samples. The STED or 4Pi-microscope setup and alignment requires relatively high light doses and sophisticated optical engineering, however providing excellent super-resolution in 2D or 3D. One technique that is widely used due to its advantageous cost-benefit ratio is dSTORM. The conceptual approach of dSTORM is analogous to the approach of STORM [2], however simplified without the need of an activator molecule. In dSTORM, a super-resolved image is obtained by subsequent imaging of well-separated subsets of blinking fluorophores. The localization of a single fluorophores can be determined with a precision exceeding the Abbe limit by far, so that dSTORM pushes the resolution to typically 30 nm without further tweaks and tricks.

For localization based methods like dSTORM the localization precision is the crucial limit for the accessible resolution which can be estimated based on the image and signal parameters ([11]– [13]) and mainly depends on signal brightness and contrast.

There have been attempts to increase super-resolution performance by maximizing the photon yield with optimized dyes [14], additives [15] or by cooling the sample down to very low temperatures [16] or doubling detection efficiency with a 4Pi-Setup [17]. Still all this approaches lack the property of being universally applicable and partly have issues regarding biocompatibility. They have therefore not reached widespread application.

Combining optical with plasmonic approaches opens exciting perspectives: So called surface plasmons in specially designed nanostructures can generate extremely high photon densities in a nanoscopic volume that is much lower than the Abbe criteria usually allows [18]. This phenomenon of metal-fluorophore interaction has been already theoretically described in 1978 [19] and a first experimental example was given by Drexhage [20]. The interaction of fluorophores with plasmonic surfaces enables amplified fluorescence, increased photostability [21] and distance dependent dynamical [22] and spectral emission shifts [23].

Several applications in the field of plasmon enhanced fluorescence imaging and spectroscopy based on metal films ([24],[25]), hyperbolic metamaterials [26] and nanoantenna structures ([27]– [31]) have already been presented. Many of the plasmonic structures mentioned here rely on a very complex nanostructure design and yield limited biocompatibility, limited applicability and sophisticated implementation.

Here we show how to use relatively simple biocompatible nanolayers to be applied in super-resolution bioimaging. A first attempt to combine the effects of metal-fluorophore interaction with super-resolution microscopy has been made by Berndt et al. [32]. Berndt and co-workers were able to determine the axial position of immobilized microtubules with nanometer precision based on the distance-dependent fluorescence lifetime modulation near a metal-coated surface. In a next step this method was also used to resolve the height profile of the membrane of a living cell with nanometer accuracy ([22],[33]). As stand-alone techniques, both approaches are only able to provide super-resolution capabilities in the axial dimension. The strength of the concept presented here lies in the straightforward implementation.

Here, we present how simple, biocompatible metal-dielectric nanocoatings can enhance the lateral resolution performance of dSTORM in a standard epifluorescence dSTORM setup. First, we characterized the technique by using the well-known geometry of fluorescently labeled microtubules. To demonstrate the potential of our optoplasmonic approach we resolved the eight-fold symmetry of the GP210-ring associated with the nuclear pore complex and show the enhanced localization precision. The nuclear pore complex is a membrane protein which is of interest to many biological questions as it plays a key role in the regulation of molecular traffic between the cytoplasm and the nucleus [34]. The eightfold symmetry of the GP210 protein associated with anchoring the nuclear pore complex in the nuclear membrane [35] can be resolved by dSTORM or STED as shown previously ([36]–[38]), however, it remains inaccessible by diffraction limited fluorescence imaging methods.

## Results

To find the ideal design of the metal-dielectric coating for our application we performed simulations of the distance dependent fluorescence enhancement for different substrate geometries considering the specific fluorophore excitation and emission properties. Figure 1a) shows a scheme of the simulation geometry. As substrates for the metal-dielectric coating, we chose conventional microscopy glass coverslips. The metal-dielectric coating consists of a silver (Ag) layer of a thickness *d*_*m*_ and silicon nitride (Si_3_N_4_) cover of a thickness *d*_*d*_. For the excitation enhancement, we compared a plane wave excitation with a wavelength of 640 nm encountering the metal-dielectric substrate with the excitation field in case of a bare glass substrate. To theoretically predict the emission enhancement, we analyzed the ratio of the distance dependent radiative rate of a dipole located at a height *h* in vicinity of a metal-dielectric substrate or a glass coverslip. The dipole excitation and emission spectra correspond to the spectra of Alexa Fluor 647 dye molecules, which are comparable to the properties of HiLyte Fluor 647 dye molecules. Based on the retrieved height dependent excitation and radiative enhancement rates we calculated an efficient enhancement rate as the arithmetic product of both. Figure 1b) shows the simulation results of excitation (dashed lines), radiation (dash-dotted lines) and fluorescence enhancement (solid lines) for two different substrate designs. The resulting enhancement curves show very distinct patterns for the two different designs as the amplitude and height distribution of the fluorescence enhancement can be tailored by the metal-dielectric substrate design. This way the maximum of the fluorescence enhancement can be placed at the height of a structure of interest for a given fluorescent label.

**Figure 1:**
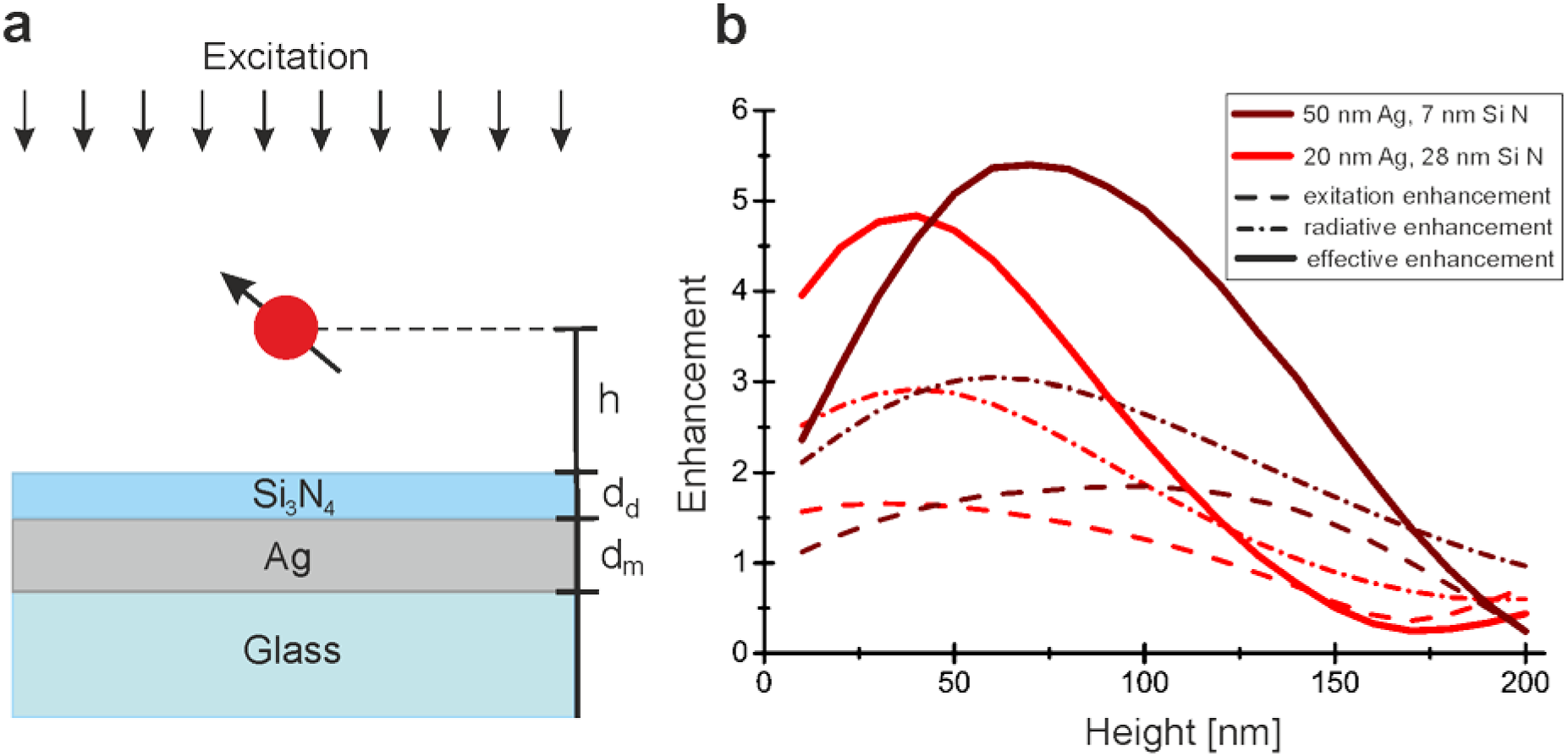
Simulation of the fluorescence enhancement on metal-dielectric substrates: **(a)** Scheme showing the geometry used for the simulations: The metal-dielectric substrate consists of glass coverslip with a silver (Ag) coating of the thickness d_m_ and a silicon nitride (Si_3_N_4_) capping layer with a thickness of d_d_. A dipole situated at a variable height h above the surface experiences a plane wave excitation. **(b)** Simulated height dependent excitation (dashed line), radiative (dash-dotted line) and effective (solid line) enhancement for a dipole with the excitation and emission properties of an Alexa Fluor 647 dye molecule.

To demonstrate our approach, we first designed and fabricated (see Materials and Methods section) a substrate, which places the maximum of the fluorescence enhancement at the height of microtubules which are labeled with HiLyte Fluor 647. Microtubules have a defined cylindrical shape with a diameter of 25 nm and can reach a length of up to 50 µm [39]. Avidin supported immobilization ensured that the microtubules are placed equally at a known distance of 7.5 ± 1 nm to the surface [32] (see Figure 2a)). With a metal-dielectric substrate composition of 20 nm silver and 28 nm Si_3_N_4_ the maximum of the emission enhancement is placed close to the height of an avidin immobilized HiLyte647-labeled microtubule. Considering the height distribution of the fluorescent markers due to the cylindrical geometry of the microtubule simulations suggest an average fluorescence enhancement of factor 4.38 ± 0.23 resulting in an estimated precision enhancement of a factor 2.

**Figure 2:**
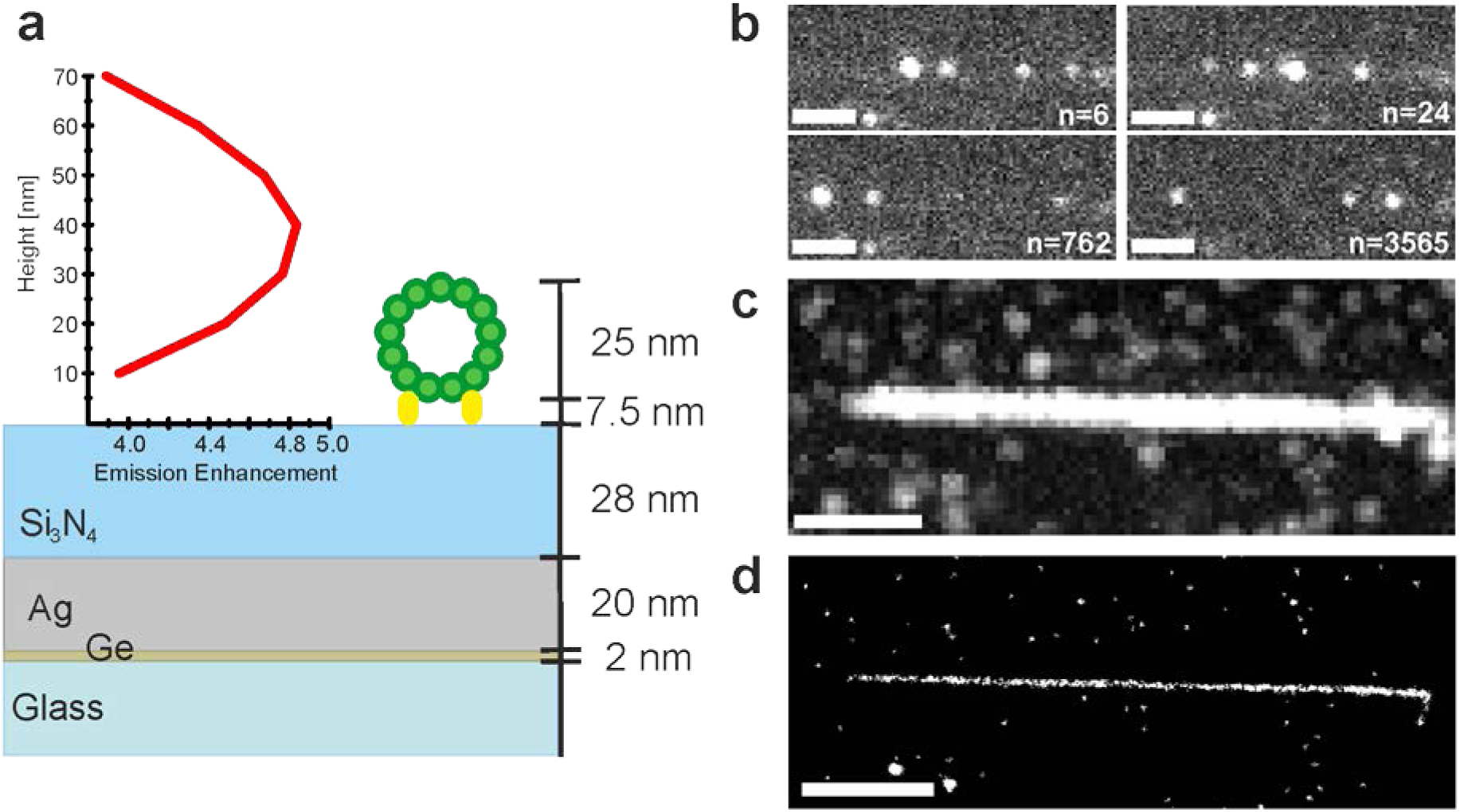
Experimental design and dSTORM experiment: **(a)** Metal-dielectric substrate design consisting of a glass cover slip with a 2 nm germanium adhesion layer, 12 nm silver layer and a 28 nm silicon nitrite layer places the maximum of the enhancement field close to the axial position of a HiLyte647-labeled microtubule. **(b)** Maximum intensity projection of the dSTORM stack of a microtubule immobilized on the substrate. **(c)** Super-resolved reconstructed average shifted histogram with an of a factor 10 magnification. Scale bars: 2 µm

To test the super-resolution performance, we carried out dSTORM experiments both on the especially designed substrate and on a bare glass coverslip under otherwise equal conditions. Both image stacks were analyzed with the same settings for localization and image reconstruction. We detected a clear fluorescence signal enhancement of a factor 1.6 that was also translated into a localization precision enhancement of a factor 1.25 (see figure 3).

**Figure 3:**
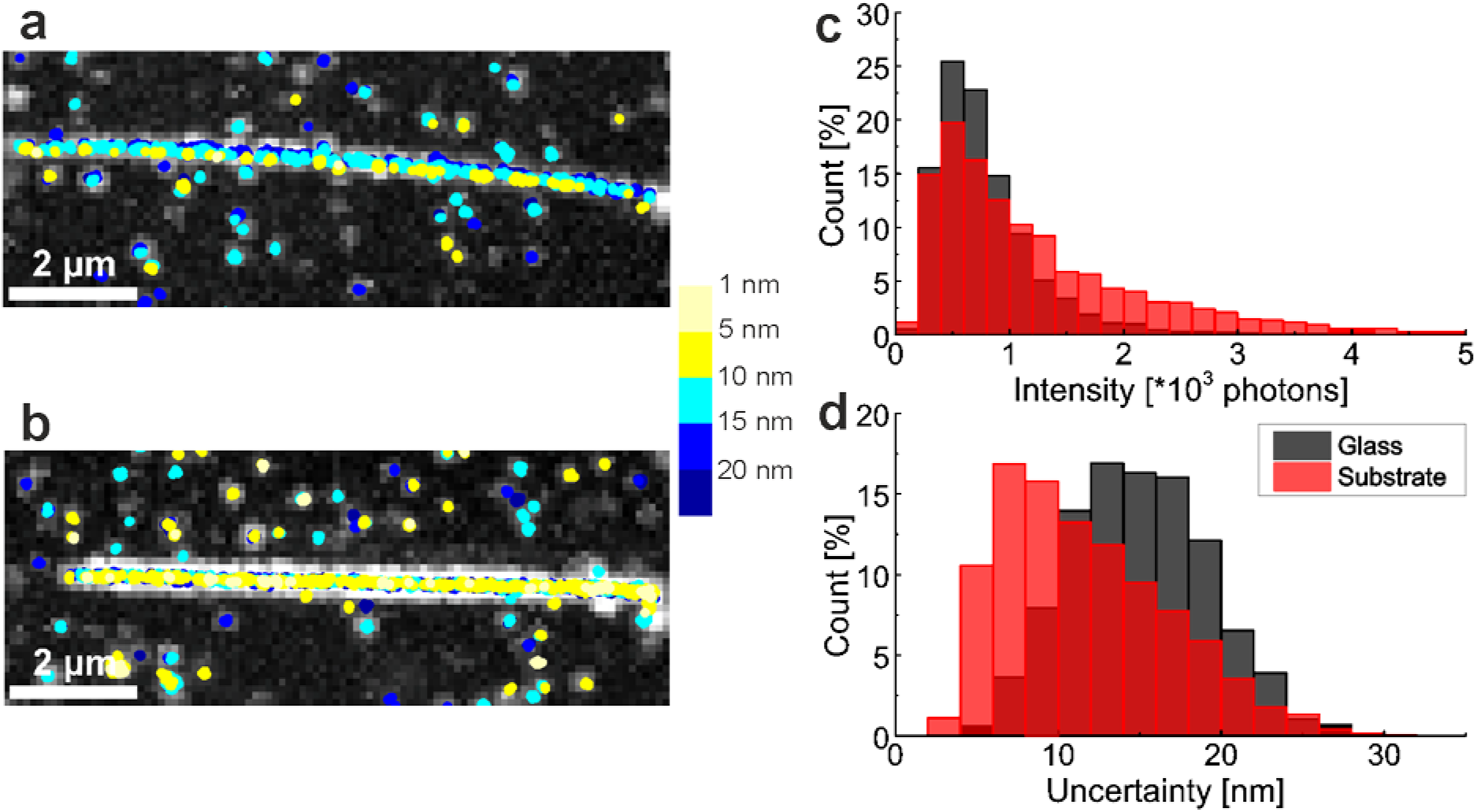
Localization microscopy on a glass coverslip and on a metal-dielectric substrate: (a+b) Overlay of scatter plot of localization positions onto maximum intensity projection with a color code indicating the related localization uncertainty of each event for a microtubule immobilized on glass coverslip (a) and on a metal-dielectric substrate (b, 20 nm Ag & 28 nm Si_3_N_4_). **(c+d)** Histogram of the intensity distribution (a) and the uncertainty distribution of the localization events for the experiment on a pure glass coverslip (grey) and the metal-dielectric coated coverslip (red).

The maximum height at which an enhancement effect can be realized with this method strongly depends on the excitation and emission wavelength range of the used fluorescent marker. To reach this height we chose a thickness of 7 nm for the Si_3_N_4_ layer to still ensure an intact dielectric spacer layer and a 50 nm thick silver layer. As simulations show, for a dye situated in the red visible wavelength regime like HiLyte647 or Alexa Fluor 647 the maximum of the fluorescence enhancement is reached at a height of around 70 nm and reaches up to around 160 nm where it has decreased to an enhancement factor of two (see figure 4). This height range is ideal for the investigation of proteins integrated into or attached to the basal membrane of an adherent cell.

**Figure 4:**
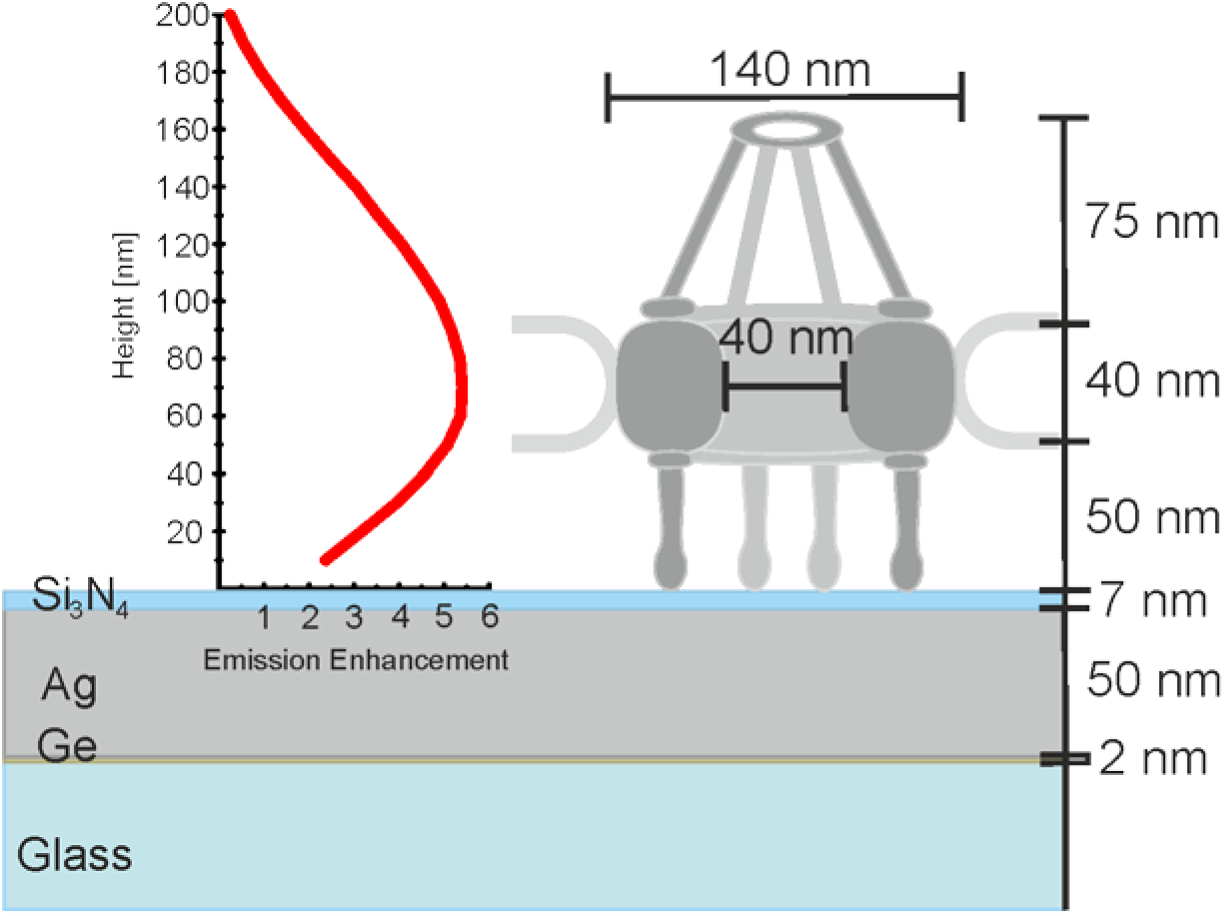
Experimental design to resolve the NPC: Metal-dielectric substrate design places the maximum of the enhancement field for an Alexa Fluor 647 dye close to the axial position of the ring structure of the nuclear pore complex. The GP210 protein located in tis ring is targeted via immunolabeling.

Cellular structures such as the nuclear pore complex cannot be resolved with conventional confocal or wide-field fluorescence microscopy which is illustrated by simulated fluorescence data (figure 5a)). Assuming a dSTORM experiment with a resolution of ∼ 15 nm the single elements of the ring appear (figure 5b) left) in the simulation. Assuming a fluorescence enhancement factor 1.6, as experimentally demonstrated in figure 3, the resolution could be further improved so that the single elements of the eightfold symmetry ring are clearly discernible (figure 5b) right). To experimentally confirm the simulation results, we performed dSTORM experiments with nuclear envelopes spread on a metal-dielectric substrate (2 nm Ge, 50 nm Ag & 7 nm Si_3_N_4_) with a control sample on a bare glass coverslip. In figure 5 c) the reconstructed super-resolved image of GP210 protein rings labelled via a X222 antibody with an Alexa Fluor 647 conjugated secondary antibody is shown for an experiment on a glass coverslip (left) and on the metal-dielectric substrate (right). We see the expected enhancement by the metal-dielectric coating resulting in an enhanced dSTORM image. The single elements of the GP210 ring can be clearly distinguished. Overall, the resolution of this image is enhanced in comparison to the situation on the bare glass coverslip due to a contrast enhancement of about a factor 2.8, which should result in a resolution enhancement of a factor 1.6.

**Figure 5:**
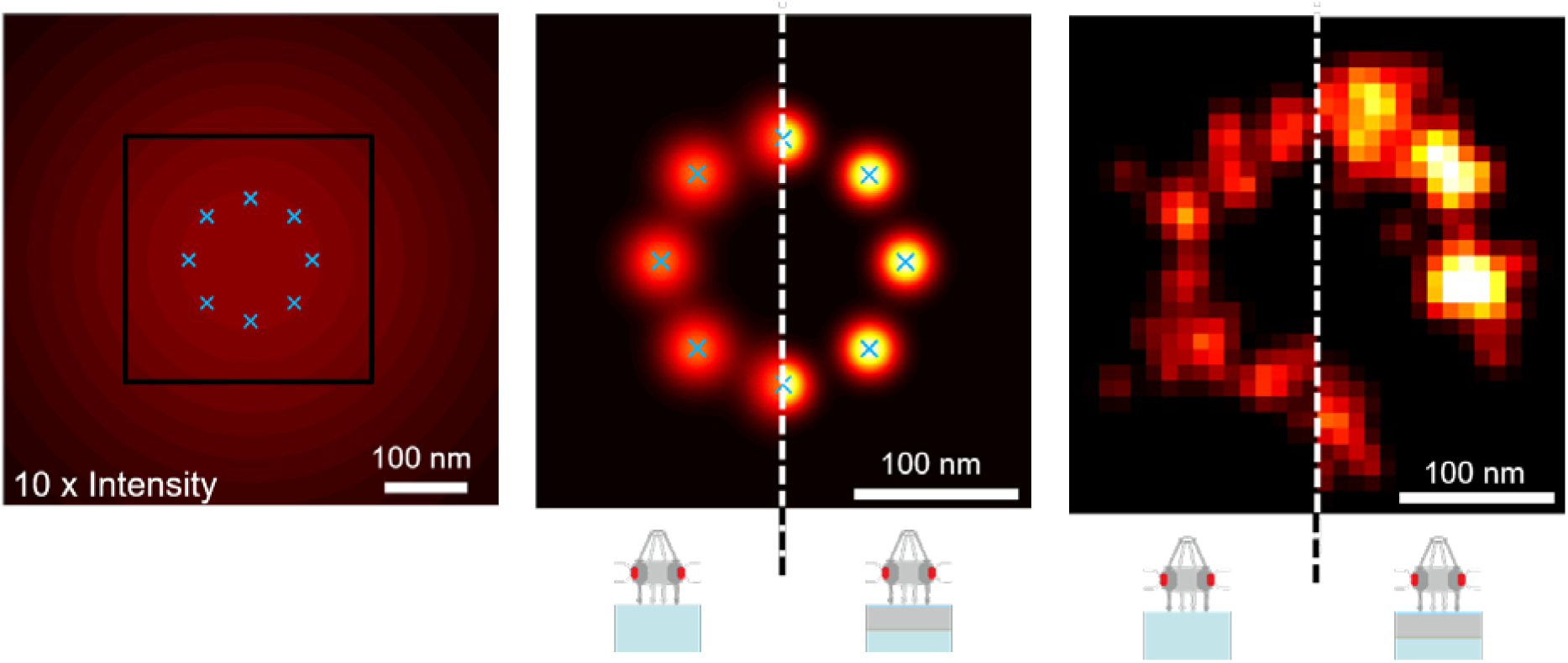
Resolving the nuclear pore complex: **(a)** Simulated diffraction limited fluorescence image of a nuclear pore complex, maximum of color scale at 10 %. **(b)** Simulated super-resolved image of the nuclear pore complex as it would be achieved by dSTORM (left) and as it could be achieved by enhanced dSTORM on a metal-dielectric substrate (right). **(c)** Experimental super-resolved images of the nuclear pore complex acquired on a glass coverslip (left) and on a metal-dielectric substrate (right)

## Discussion

In conclusion, we have shown that by an anticipatory design of silver-silicon nitride substrates the localization precision of dSTORM is enhanced when imaging microtubules or the nuclear pore complex. The metal-dielectric coated substrates are a versatile tool: The simple tree-ply design of our metal-dielectric coatings grants a straightforward one-step fabrication and allows tailoring the shape of the resulting enhancement field to the sample geometry while taking into account the excitation and emission properties of the fluorescent label at hand. Besides this, the silicon nitride-coating prevents fluorescence quenching and assures the biocompatibility of the substrate allowing the cultivation of living cells. Here we fabricated the described nanostructures under clean-room conditions. However, those could be even fabricated by non-specialists and without clean-room access using suitable tabletop thin film deposition systems, which will be implemented by us in future, work.

The handling of the prefabricated metal-dielectric substrates is user-friendly, as it does not require any further training or caution. Thus, the technique provides a high degree of versatility concerning application and implementation in any (dSTORM) microscopic setup where only the coverslip is modified. As other optoplasmonic approaches our method is surface bound, which only takes effect up to a height of about 160 nm above the substrate interface. Here, this is beneficial as allows for selectively enhancing the fluorescence signal in the plane right in vicinity above the surface while suppressing background signal from regions further away. This is increasing the contrast like in an evanescent filed illumination approach as for total internal reflection fluorescence (TIRF). We understand that this could also be a limitation in other applications resulting in limited or insufficient penetration depth. In principle, it is possible to extend the axial range of the enhancement field when exciting long range plasmons. But as the confinement of the field decreases with growing surface distance the beneficial enhancement effects would be diminished when spread over a larger operating range.

As we have demonstrated, this approach is especially interesting for super-resolution imaging, but it could also be advantageous in contrast enhanced single molecule tracking or single molecule detection in high throughput screening assays.

The simulations even predict a signal enhancement of up to a factor four, which would translate into a precision enhancement of a factor two. The maximum values were not yet reached experimentally, most likely due to underestimation of the sample’s distance or photophysical effects we are not (yet) aware of. We believe, even with the current enhancement, this technique can tip the scales in favor of a clearly resolved structure with further experimental developments of such an optoplasmonic approach becoming more than promising.

## Acknowledgements

We thank Jörg Enderlein and Rainer Heintzmann for fruitful discussions and Lena Lauber for assistance with the dSTORM experiments. This work was financially supported by the German Research Foundation (DFG) (SFB/TR 166: K.G.H., H.S.H, M.S.), the Rudolf Virchow Center of the University of Würzburg, the IDK Receptor Dynamics of the Elite Netzwerk Bayern (B.S.), the University of Würzburg (M.E., M.K., S.H.) and the State of Bavaria (clean room facilities).

## Author Contributions

K.H., M.S. and H.S.H conceived and designed the experiments. H.S.H. and B.S. performed the simulations and H.S.H performed experiments and analyzed the data. M.E. fabricated the metal-dielectric coatings. S.H. supervised the fabrication process. M.K. advised the first nanofabrication steps and provided quality control for the final metal-dielectric substrates. K.H. and H.S.H wrote the paper.

## Author Information

The authors declare no competing financial interests. Correspondence and requests for materials should be addressed to K.H.

## Methods

### Substrate fabrication

The metal-dielectric substrates were fabricated on 170 µm thick glass coverslips (Menzel Coverslip 24×24 mm, #1.5, selected, Fisher Scientific, Schwerte, Germany), which were cleaned in piranha-solution (3 parts of 95-98 %H_2_SO_4_ (Roth, X944) and 1 part of 30 % H_2_O_2_ (AppliChem, A0626)) at 70 °C for 20 minutes and dried at 100 °C for 10 minutes prior to fabrication. A 2 nm germanium wetting layer was deposited by electron beam evaporation followed by a 20 nm or 50 nm thick sliver layer. Finally, a dielectric spacer layer was applied on top by RF- sputtering of silicon nitrite until a thickness of 23 nm or 7 nm was reached. The metal-dielectric substrates were stored under vacuum.

### Microtubule sample preparation

20 µg of lyophilized HiLyte 647 labeled porcine tubulin (tebu-bio, 027TL670M-A) were diluted in 5 µl of the ice-cold polymerization buffer (1 mM GTP (tebubio, BST06), 5 mM MgCl_2_ (AppliChem, A1036) and 5 % DMSO (AppliChem, A1584) in BRB80) and placed on ice for 5 minutes before incubating the tubulin solution at 37 °C for 2 hours. The microtubules were stabilized by adding 195 µl of 10 µM paclitaxel (tebu-bio, TXD01) in BRB80. After spinning down the microtubules at 16100 × g and 23 °C for 20 minutes the supernatant was discarded and the remaining pellet was resuspended in 200 µl of 10 µM paclitaxel in BRB80. The polymerized and stabilized microtubules were stored at room temperature and used within the next week. Before an experiment, a flow cell was constructed by placing two 2 mm stripes of double sided tape (TESA) along the long axis of a 170 µm thick 24x40 mm glass coverslip (Menzel Coverslip 24×40 mm, #1.5, selected, Fisher Scientific, Schwerte, Germany) and placing a plane 24×24 mm coverslip or a metal-dielectric substrate (20 nm silver and 23 nm silicon nitrite) with the coating facing inward on top. Prior to this step, the plane glass coverslips were cleaned by sonicating for 1 hour in chloroform (Sigma, 472476), drying, sonicating for 1 hour in 5 M NaOH (Roth, 6771) solution and washing in ddH_2_O. The cleaned glass coverslips were stored in ethanol (Sigma, 32205). The metal-dielectric substrate was washed in ethanol and dried with N_2_ just before the flow cell construction. For the immobilization of the microtubules on the surface, the microtubule solution was diluted 1: 10 in 10 µM paclitaxel in BRB80 and centrifuged for 20 minutes at 16100 × g and 23 °C. The remaining pellet was resuspended in 10 µM paclitaxel in BRB80. 20µl of the resulting purified microtubule solution was incubated for 10 min in the flow channel. After a 40 µl wash with 10 µM paclitaxel in BRB80, 20 µl of a 1 µM avidin solution (Sigma, A9275) with 10 µM paclitaxel were incubated for 5 minutes in the channel before washing again with 40 µl of 10 µM paclitaxel in BRB80. Finally, the microtubules were fixed with 2 % glutaraldehyde (Merck, 104239) and 10 µM paclitaxel solution for 10 minutes and stored in BRB80.

### Nuclear pore complex sample preparation and labeling

Nuclear envelopes were manually isolated and spread on cover glasses or substrates as described elsewhere ([36]). After a 15 minute fixation step in 3 % formaldehyde (Sigma, F8775) in PBS (Sigma, D1408) the nuclear envelopes were washed in PBS and blocked for 20 minutes in 5 % BSA (Sigma, A3983) in PBS. For the immunolabeling the nuclear envelopes were incubated for 1 hour with X222 antibodies (provided by Georg Krohne [40]), washed for 10 min in PBS and incubated another hour with Alexa647 F(ab’)_2_ fragments of goat anti-mouse IgG (Life Technologies GmbH, A21237). After a final PBS washing step for 20 minutes the nuclear envelopes were incubated in 3 % formaldehyde in PBS for 10 minutes and stored in PBS. Before the experiment, a flow cell was constructed as described in the previous section by placing the glass coverslip or substrate carrying the nuclear envelopes on top with the envelopes facing inside.

### dSTORM imaging

The dSTORM imaging was performed with an inverted light microscope (Zeiss Observer Z.1, Carl Zeiss AG) equipped with an EMCCD camera (iXon ultra DU-897U-CSO-#BV, Andor, Belfast, UK), a C-APOCHROMAT 1.15 NA, x63 water objective (LD C-Apochromat 63x NA 1.15, 421887-9970, Carl Zeiss AG) and an optovar with 2.5-fold magnification. The excitation was performed with a 640 nm laser (iBEAM-smart-640-S, Toptica) through an illumination lens with 40 mm focal length, resulting in an effective illumination intensity of 1-5 kW/cm^2^. In the excitation light path a clean-up filter (640/8 nm MaxDiode^™^ laser clean-up, Semrock) is passed before a dichroic mirror (BrightLine quadedge 405/488/532/635, Semrock) separates excitation and emission light. In the detection path an additional notch filter (StopLine quadnotch ZET 405/488/561/647, Semrock) and a bandpass filter (700/75 ET, Chroma) are placed. After applying the dSTORM buffer (100 mM or 125 mM MEA (Sigma, M6500), 20 mM D-glucose (Sigma, G7528), 0.55 mg/ml gluco-oxidase (Roth, 60281), 0.011 mg/ml catalase (Sigma, C1345) in PBS adjusted to pH 7.7 with 1M KOH (Sigma, 30603) solution) image stacks of 20000 frames were acquired with an exposure time of 5 or 10 ms.

### Image reconstruction and analysis

dSTORM images were reconstructed with the ImageJ plugin ThunderSTORM [41]. For each experiment, the same localization parameters were used. For the microtubule experiments, localizations with less than four neighbors in a 60 nm radius were discarded. In case of the nuclear pore complex, reconstruction localization duplicates were filtered by merging localization events that repeatedly occurred within a radius of 15 nm with a maximum of 50 off-frames in between. Only localizations with an intensity of more than 500 and less than 5000 photons were considered and in order to discard events from unspecific background a density filter of a minimum of 5 events in a radius of 50 nm was applied.

### Simulation

Simulations of the enhancement effect were performed with the commercial software Comsol Multiphysics^™^ 4.4. For the excitation enhancement, a plane wave excitation encountering the metal-dielectric substrate of the experimentally used geometry was compared with the excitation field in case of a plane glass substrate. In order to theoretically predict the emission enhancement, the distance dependent radiative rate of a dipole located near a metal-dielectric substrate of a glass coverslip was compared. Based on the retrieved height dependent excitation and radiative enhancement rates an efficient enhancement rate was calculated. In case of the microtubules the height distribution of the fluorophores is very well defined by the sample geometry and was also considered. Simulations of the fluorescence images of the nuclear pore complex were calculated with MatLab (Mathworks Inc., Natick, MA).

